# Hippocampal theta and episodic memory

**DOI:** 10.1101/2022.03.13.484014

**Authors:** Joseph H. Rudoler, Nora A. Herweg, Michael J. Kahana

**Affiliations:** Computational Memory Lab, Department of Psychology, University of Pennsylvania, PA, USA 19104; Department of Neuropsychology, Institute of Cognitive Neuroscience, Faculty of Psychology, Ruhr University Bochum, Germany 44780

## Abstract

Computational models of rodent physiology implicate hippocampal theta as a key modulator of learning and memory (Buzsaki & Moser, 2013; J. E. Lisman & Jensen, 2013), yet human hippocampal recordings have shown divergent theta correlates of memory formation. Herweg et al. (2020) suggest that decreases in memory-related broadband power mask narrowband theta increases. Their survey also notes that theta’s role in memory appears strongest in contrasts that isolate retrieval processes and when aggregating signals across large brain regions. We evaluate these hypotheses by analyzing human hippocampal recordings captured as 162 neurosurgical patients (*N* = 86 female) performed a free recall task. Using the irregular-resampling auto-spectral analysis to separate broad and narrow-band components of the field potential we show: 1) Broadband and narrowband components of theta exhibit opposite effects, with broadband signals decreasing and narrow-band theta increasing during successful encoding; 2) Whereas low-frequency theta oscillations increase prior to successful recall, higher-frequency theta and alpha oscillations decrease, masking theta’s positive effect when aggregating across the full band; 3) Theta’s effects on memory encoding and retrieval do not differ between reference schemes that accentuate local signals (bipolar) and those that aggregate across large reference (whole brain average). In line with computational models that ascribe a fundamental role for hippocampal theta in memory, our large-scale study of human hippocampal recordings shows that 3-4 Hz theta oscillations reliably increase during successful memory encoding and prior to spontaneous recall of previously studied items.

**Significance statement:** Analyzing recordings from 162 patients we resolve a long-standing question regarding the role of hippocampal theta oscillations in the formation and retrieval of episodic memories. We show that broadband spectral changes confound estimates of narrowband theta activity, thereby accounting for inconsistent results in the literature. After accounting for broadband effects, we find that increased theta activity marks successful encoding and retrieval of episodic memories, supporting rodent models that ascribe a key role for hippocampal theta in memory function.

## 1 Introduction

Since the classic work of (Scoville & Milner, 1957), we have known that the hippocampal formation plays a crucial role in human context-dependent (episodic) memory. Whereas lesion studies reified the single-case study of H.M. (Squire, Knowlton, & Musen, 1993), further advances in our understanding of hippocampal physiology arose from recording field potentials and neuronal spiking in awake behaving mammals (e.g., O’Keefe & Dostrovsky, 1971; Knierim, Kudrimoti, & McNaughton, 1995; McNaughton, Barnes, & O’Keefe, 1983). These studies led to discoveries regarding the role of theta oscillations and place cell activity in animal learning (see J. Lisman, Jensen, & Kahana, 2001, for a review). While human scalp EEG studies had suggested some role for theta rhythms in cognitive processes (e.g., Schacter, 1977) it was only at the turn of the 21st century that depth-electrode recordings in neurosurgical patients specifically implicated theta oscillations in human spatial and verbal memory (Kahana, Sekuler, Caplan, Kirschen, & Madsen, 1999; Sederberg, Kahana, Howard, Donner, & Madsen, 2003; Ekstrom et al., 2005). The ability to record neural activity from indwelling electrodes synchronized with computer-controlled memory experiments spawned a series of important discoveries regarding the electrophysiology of human learning and memory (Johnson & Knight, 2015)

Despite recent progress in the neurophysiology of human memory, considerable confusion surrounds the role of hippocampal theta activity in key memory processes, such as successful encoding and retrieval. To isolate neural correlates of successful memory encoding, researchers typically sort studied items into two groups: those that are subsequently recalled or recognized and those that are subsequently “forgotten”. Neuroimaging studies employing this contrast have frequently identified the hippocampus as a region of increased haemodynamic activity during successful encoding. To isolate neural correlates of memory retrieval, researchers often compare the period during which recollection occurs with a control period comprising either a matched deliberation interval (Burke, Sharan, et al., 2014) or a period preceding a retrieval error (Long et al., 2017).

In a recent review Herweg et al. (2020) identify a highly inconsistent pattern of findings, particularly with regards to data from direct recordings from the human medial-temporal lobe (MTL). They find that most studies either report negative associations between MTL theta and memory, or mixed patterns of results with some electrodes exhibiting increases and others exhibiting decreases in theta power. Herweg et al (2020) propose several possible accounts for the discrepancies across these studies. First, they suggest that estimates of theta-band spectral power are confounded with broadband power changes, with the former reflecting synchronous oscillations and the latter reflecting asynchronous broadband activity indicative of greater attentiveness or cognitive engagement (Burke, Ramayya, & Kahana, 2015; Hanslmayr, Staudigl, & Fellner, 2012; Miller et al., 2014; Voytek & Knight, 2015). They claim that the standard subsequent memory effect (SME) analysis will emphasize changes in global attention rather than memory-specific encoding processes, and suggest that negative effects largely reflect broadband activity which masks the positive theta effects in the data. Theta oscillations might emerge after making analytic corrections for broadband changes. They also note that studies looking at retrieval processes - which are not bound to external stimuli and compare more similar attentive states - may be less susceptible to broadband confounds and therefore more suited to identify increases in hippocampal theta. Memory retrieval in a free recall task also depends critically on associative processes that bind items together based on a combination of their temporal and semantic relations (Kahana, 2020), which may themselves be linked to the strength of theta oscillations (Solomon, Lega, Sperling, & Kahana, 2019). Finally, they note that most magnetoencephalography (MEG) and scalp EEG studies find positive theta effects, often overlying frontal regions. While both invasive and non-invasive studies have yielded mixed results, the noticeably greater proportion of non-invasive studies reporting positive theta effects raises questions about how theta is impacted by the spatial resolution of the recording technology and the referencing scheme applied to the signal. It may be that bipolar reference schemes used in many of the studies reporting decreases in theta may have filtered out theta increases that appear synchronously over larger brain areas.

The present paper evaluates these accounts of the discrepant findings concerning hippocampal theta and memory. To do so, we analyze a large dataset comprising 162 neurosurgical patients fitted with hippocampal depth electrodes. Using standard wavelet methods we analyze spectral activity during encoding and retrieval phases of a delayed free recall task. We also separate broadband and narrowband components of spectral activity using irregular-resampling auto-spectral analysis (IRASA) (Wen & Liu, 2016). To evaluate the hypothesized role of hippocampal theta in memory encoding, we analyze the period during which an item is studied and compare trials for which the stimulus is subsequently recalled or forgotten. At retrieval, we compare the period immediately preceding verbal recall with matched periods of deliberation, when patients are trying to recall but no items come to mind. Finally, we evaluate the hypothesis that local spatial referencing obscures theta’s role in memory by repeating the above comparisons separately using a global average reference of implanted electrodes (as compared with a bipolar reference that localizes activity to the differential voltage between neighboring channels).

## 2 Materials and Methods

### 2.1 Subjects

We analyzed hippocampal depth-electrode recordings from 162 patients who participated in the DARPA-funded *Restoring Active Memory* project (e.g., Ezzyat et al., 2018; Solomon, Stein, et al., 2019; Solomon, Lega, et al., 2019; Phan, Wachter, Solomon, & Kahana, 2019). This publicly-shared dataset includes >300 patients with drug-resistant epilepsy who took part in memory testing while undergoing a neurosurgical procedure to localize seizure activity and functional tissue. These patients (*N* = 86 female, *N* = 17 left-handed) had a mean age of 38 (ranging from 18 to 64). Researchers obtained informed consent from each patient and the research protocol was approved by the institutional review board (IRB) at the University of Pennsylvania and each participating hospital.

Patients contributed variable numbers of trials (i.e. studied test lists) depending on the length of their hospitalization and their interest in participating. We analyzed data only from patients who recalled at least, on average, one word per list; we also limited this study to patients with at least one bipolar pair consisting of contacts localized within the hippocampus (see *Electrode Recording: Localization and Preprocessing*).

### 2.2 Experimental Design

Patients completed a free recall task, in which they encoded a sequence of 12 words which appeared on a blank screen for 1600 ms each during a study phase. The spacing between words is jittered between 750 and 1000 ms. Following each study phase patients performed a 20 s arithmetic distractor task in which they solved problems of the form *X* + *Y* + *Z* =??, where *X*, *Y*, and *Z* were positive or negative numbers between 1 and 9. Responses were made on a keypad, with presentation of additional math problems following each response (i.e., a self-paced task). After the delay, a row of asterisks accompanied by an 800 Hz auditory tone signaled the start of the recall period. At this point patients recalled out loud all the words they could remember from the list in 30 seconds. They repeat this sequence 25 times to complete the experiment, but not all patients complete a full 25 trials. Many patients repeat this process for multiple experimental sessions.

### 2.3 Electrode Recordings: Localization and Preprocessing

Our study focuses specifically on neural recordings from the hippocampus, defined as including regions CA1, CA2, CA3, CA4, dentate gyrus, and subiculum. To localize the recording contacts of depth electrodes, first hippocampal subfields and MTL cortices were automatically labeled in a pre-implant, T2-weighted MRI using the automatic segmentation of hippocampal subfields (ASHS) multi-atlas segmentation method (Yushkevich et al., 2015). Next, post-implant CT images were manually annotated with the voxel coordinates of individual recording contacts in CT-space. Post-implant CT images were coregistered with presurgical T1 and T2 weighted structural scans with Advanced Normalization Tools (Avants, Epstein, Grossman, & Gee, 2008), aligning the locations of individual recording sites to the anatomical labels assigned through automatic segmentation. For the majority of subjects in this dataset, MTL depth electrodes that were visible on overlaid CT and MRI scans were then manually annotated with localizations within MTL subregions by neuroradiologists with expertise in MTL anatomy. The electrodes used in this analysis appear in Figure 1 in transformed MNI coordinate space. Algorithms that perform automatic segmentation and coregistration naturally introduce some imprecision, especially for patients with lesions or otherwise altered anatomy. Moreover, surgically implanted depth electrodes displace brain tissue and further complicate this task. Our confidence in these localizations stems from the manual work of the research team in visually inspecting the alignments and segmentations for every patient, and from the work of expert neuroradiologists checking the veracity of the anatomical labels.

**Figure 1:**
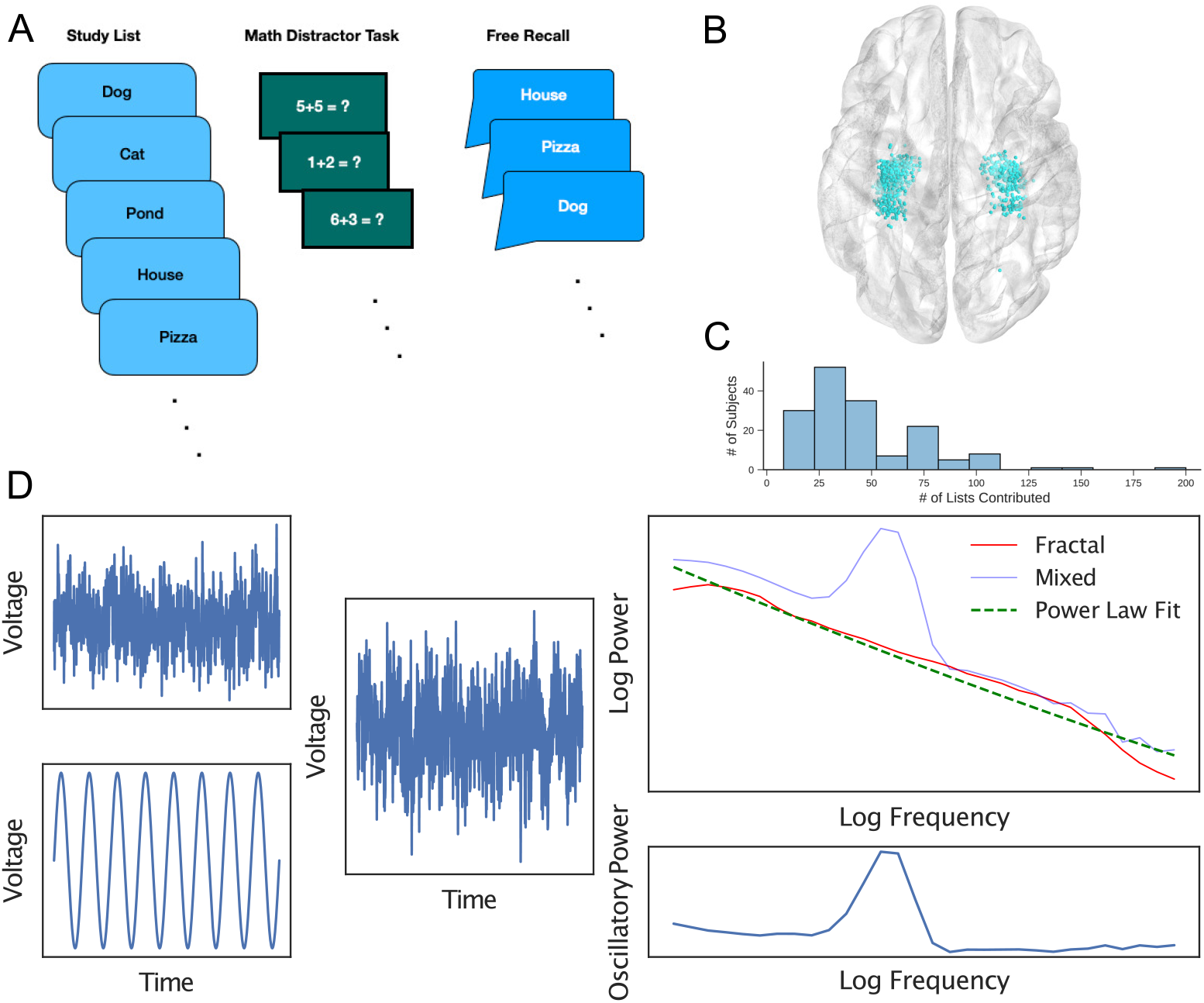
Methods. **A)** Patients performed a free recall task in which they were presented with a list of words on a blank screen in sequence, completed a math distractor task, and were then prompted to recall as many of the presented words as possible during a 30 second free recall period. **B)** Data for this analysis were collected from electrodes located in the bilateral hippocampus across 162 patients. **C)** Distribution of the number of studied lists across patients **D)** IRASA treats an EEG trace as a linear combination of an oscillatory component and a “fractal” pink noise component which is assumed to follow a power law distribution. IRASA capitalizes on a mathematical property of fractals called *self-affinity* that causes them to behave differently under re-sampling than other signals, thereby allowing us to separate the components and obtain a purely oscillatory power spectrum.

The original sampling rates for these recordings vary by hospital and patient, but are all at least 500 Hz. For analysis, we resampled each recording to 500 Hz for consistency. We re-referenced the EEG using a bipolar montage in order to mitigate noise and increase the spatial resolution of the recordings, except for one analysis that explicitly compares this bipolar reference to a global (or whole-brain) average reference scheme. Applying a Butterworth bandstop filter of order 4 removed 60 Hz line noise from the recordings.

### 2.4 Separating broadband and narrowband effects with IRASA

IRASA, introduced by Wen and Liu (2016), is a method for separating oscillations from the pink noise background. We first assume that the EEG timeseries is a mixed signal containing both fractal (*f* (*t*)) and oscillatory (*x*(*t*)) components.

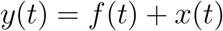

Fractals are mathematically interesting for a number of reasons, but a property of particular importance is that they exhibit *self-affinity*. This means that fractals are scale free; geometrically, magnifying a portion of a fractal will produce qualitatively the same pattern. Expressed mathematically, when a fractal time series is resampled by a factor *h*,

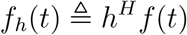

which means that the statistical distribution of the resampled time series is the same as the statistical distribution of the original time series multiplied by a scaling term (The Hurst exponent *H* is related to the time series’ auto-correlation). In the frequency domain, this self-affinity manifests even more directly as

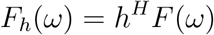

which states that the Fourier transform after resampling is equal to the Fourier transform of the original time series multiplied by a scaling factor. This property is useful because resampling causes non-fractal signals to shift in the frequency domain. For an example of how signals typically shift in frequency space under resampling, consider an oscillation at 5 Hz in a recording sampled at 1000 Hz. This oscillation completes a full cycle every 200 samples. If the recording is downsampled to 500Hz, then essentially every other sample is removed. Now, the same 5 Hz oscillation completes a cycle in only 100 samples. Without properly correcting for the change in sampling rate, it appears as if the speed of the oscillation has doubled to 10 Hz. To operationalize the property of self-affinity, we use the discrete Fourier Transform to compute the autopower spectrum of the time series up- and down-sampled by a set of non-integer factors *h*. Taking the median across the full range of h values removes the shifted oscillatory peaks, leaving exactly the fractal component. Subtracting this fractal component from the overall power spectrum then isolates oscillations. Figure 1 shows an example of the method applied to simulated data. We refer the reader to the original methods paper (Wen & Liu, 2016) for a more mathematically detailed description of IRASA.

#### 2.4.1 Isolating rhythmic oscillations

In order to isolate the oscillatory components of the neural power spectrum, we applied IRASA to an epoch of 300-1300ms following word presentation for every event in the task’s encoding phase. To study the electrophysiology of memory retrieval we repeated the analysis for the epochs from 800-50ms prior to recall vocalization. These time windows were chosen to balance the trade-off between having a sufficiently long window to assess power in the low theta band and being sufficiently specific to an individual, temporally punctate behavioral event.

IRASA decomposes the power spectrum into fractal and oscillatory components for each event and channel within every patient. The choice of resampling factors *h* controls the extent to which the method is robust to outliers, but trades robustness for spectral smoothing that decreases our frequency resolution. As we wanted to ensure our analysis did not unintentionally include noise artifacts created by large oscillations, but are also interested in having a high-resolution spectral decomposition that can distinguish between different narrowband effects, we chose a relatively conservative set of resampling factors from 1.1 to 2.0, linearly spaced by 0.05. This is the default set of resampling factors recommended and used in the original methods paper.

As IRASA explicitly extracts only the fractal component from the power spectrum, in order to isolate the oscillatory component we need to take the difference of the full spectrum and the fractal component. This is simple enough, but poses a challenge when log-transforming to suppress extreme values and normalize the data. If the fractal estimate is greater than the mixed power spectrum, the oscillatory power will be negative and the log transform undefined. We therefore introduce a Shifted Symmetric Log transform (SSL) to achieve the same goals without issue. This transform is defined as follows:

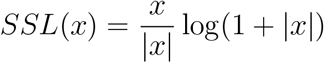

This function retains the useful properties of the logarithm, but it is symmetric about the *x*-axis and does not go to negative infinity at very small values.

#### 2.4.2 Wavelet power

We computed wavelet power at both encoding and retrieval to serve as a baseline against which we can compare our results using IRASA. We computed power at logarithmically spaced frequencies using Morlet wavelets with a width of 4 cycles. For this analysis, we included buffers on either side which corresponded to at least two cycles at the lowest frequency being analyzed (to avoid edge effects). At retrieval, we excluded recalls which were preceded by a vocalization during the buffer period preceding the epoch; we also implemented a mirrored buffer following the epoch in order to avoid contaminating our spectral estimates with vocalization artifacts from recall onset.

### 2.5 Statistical Analyses

Our primary question in analyzing item encoding is the following: how does power at a given frequency (e.g. theta) behave as a function of subsequent recall status? In answering this question we want to account for individual differences as well as session-level effects; memory performance and power might differ across patients or event across different recording sessions for the same individual. Accordingly, we fit a linear mixed effects model for each frequency of interest using the *lmer4* package in R (Bates, Mäachler, Bolker, & Walker, 2015) which predicts channel-averaged power as a function of subsequent recall (1 for success and 0 for failure), with random slopes and intercepts for the effects of subject, session, and trial (studied word list). To ensure proper estimation of the effects and their standard errors, we started with a maximal model and incrementally reduced the model complexity to remove zero-variance components and avoid singularities in the estimated variance-covariance matrix (Matuschek, Kliegl, Vasishth, Baayen, & Bates, 2017; Bates, Kliegl, Vasishth, & Baayen, 2018). Our analysis of memory retrieval followed the same procedure and model, but the binary memory success variable represented successful memory retrieval (as opposed to baseline deliberation) rather than successful memory encoding (as determined by subsequent recall).

We report effects at each frequency of interest as *β* coefficients from this model, and use a Wald test to evaluate statistical significance. We then correct for multiple comparisons by using the Benjamini-Hochberg procedure for controlling false discovery rate. This method is appropriate when tests are positively correlated, as are spectral estimates at similar frequencies.

### 2.6 Data and Code Availability

Raw electrophysiogical data used in this study are available upon request from https://memory.psych.upenn.edu/Data_Request. Analysis and data visualization code is also available for direct download from https://memory.psych.upenn.edu/files/pubs/RudoEtal22.code.tgz. The Python implementation of the IRASA method used for this study is publicly available at https://github.com/pennmem/irasa, and other custom processing scripts used for this project can be found at https://github.com/pennmem/.

## 3 Results

As our analyses sought to elucidate the role of hippocampal theta oscillations in episodic memory encoding and retrieval, we identified all patients with hippocampal electrodes in the multi-center *Restoring Active Memory* project (see Methods). Out of a total of N=281 patients who completed the same free recall task, 162 had at least one bipolar electrode pairs whose contacts both fell within the hippocampal formation defined as including regions CA1, CA2, CA3, CA4, dentate gyrus, and subiculum, but excluding non-hippocampal MTL regions such as perirhinal and parahippocampal cortices. Each patient performed a memory task in which they studied 12 words which they attempted to recall following a brief arithmetic distractor task (see Fig.1A for a schematic of the experimental task). During the 30 s recall period, patients attempted to say as many words as they could remember from the most recent list, in any order. Each patient contributed data from multiple study-test trials (see *Methods* for details). As patients performed this memory task, intraparenchymal depth electrodes captured hippocampal field potentials (Figure 1B; see *Methods* details regarding electrode localization).

The present study seeks to clarify the role of hippocampal neural oscillations in the formation and retrieval of episodic memories. As most prior work involving direct brain recordings used standard spectral decomposition procedures (e.g., wavelet transforms, multi-tapers, other windowed FFT methods, etc.) these studies cannot disambiguate oscillations from broadband components of neural activity underlying successful mnemonic function. To address this limitation, we analyzed neural signals using the irregular-resampling auto-spectral analysis (IRASA), which exploits the fractal properties of the power-law distributed broadband component to isolate it from the mixed power spectrum (see *Methods*). Figure 1c shows how IRASA decomposes a simulated EEG trace into broadband and narrowband components. The simulated data in 1c contain a single sine wave at a known frequency (“narrowband”) and pink noise (“broadband”). IRASA estimates this broadband component (see *Methods*) and subtracts it from the mixed autopower spectrum to isolate the residual oscillatory power.

The formation of episodic memories occurs when patients study words for a subsequent recall task. To identify the spectral correlates of successful encoding we examined the 1 second interval beginning 300 ms following item presentation, thereby excluding brain signals related to perceptual processing of the presented word. Spectral analyses of hippocampal field potentials during this encoding period typically show a tilt in the power spectrum, with decreases in low frequency power and increases in high frequency power predicting subsequent recall (Burke, Long, et al., 2014; Ezzyat et al., 2017; Fellner et al., 2019). This spectral tilt manifests as a flattening of the overall spectrum in log-log space, resulting in a change in the power law exponent. IRASA, by isolating the power-law distributed background and removing that component from the power spectrum, reveals a more accurate estimate of the narrowband oscillatory patterns that coexist with broadband changes.

As shown in Figure 2, the mixed autopower spectrum - analogous to traditional wavelet or multitaper methods - shows the expected theta decreases during successful encoding. Likewise, the isolated broadband component shows decreases in theta power. The oscillatory spectrum, however, shows theta increases. The estimates of effect size and significance derive from the mixed effects model described in *Methods*, which predicts power at a frequency of interest as a function of recall status, while accounting for the effects of subject and session.

**Figure 2:**
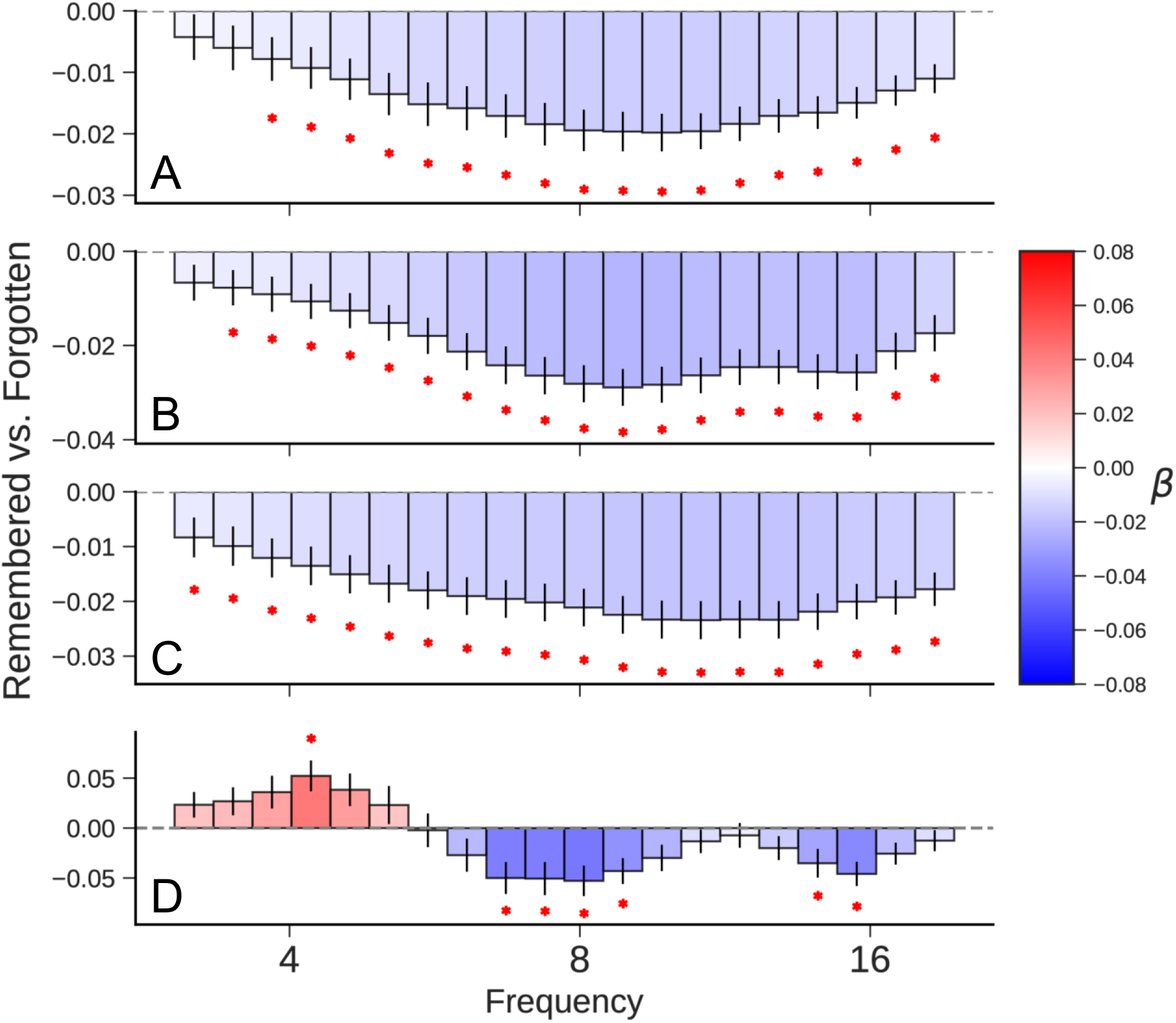
Subsequent Memory Effect. Comparison of spectral power for successful and unsuccessful memory encoding events (remembered - forgotten) based on hippocampal depth electrode recordings from 300-1300ms following item presentation. Values represent fixed effect coefficients for the effect of successful recall on spectral power. A red asterisk indicates that the effect size is significant after correcting for multiple comparisons (*p* < .05). **A)** Power computed using traditional Morlet wavelets. **B)** The mixed power spectrum (before separating broadband and narrowband effects) shows the expected theta and alpha decreases, analogous to the results using wavelets. **C)** The fractal power spectrum (broadband only) likewise shows broad decreases in low frequency. **D)** The oscillatory power spectrum, which is computed as the difference of the mixed and fractal power specra, exhibits an increase in theta power, while retaining the same decrease in alpha power.

Our finding of a small but reliable positive theta subsequent-memory effect in the human hippocampus contrasts with previous studies that found predominantly negative theta effects using wavelet methods and aggregate indices of hippocampal activity. These results confirm the hypothesis offered by Herweg et al. (2020) that broadband decreases in low frequency activity mask narrowband increases in theta activity. Given that aggregating hippocampal recordings from 162 patients only yielded a modest positive theta SME, it is likely that many studies comprising smaller samples would not be powered to detect this effect (e.g., Sederberg, Schulze-Bonhage, Madsen, Bromfield, Litt, et al., 2007). We next turn to the question of memory retrieval, asking whether isolation of narrow-band spectral components can resolve mixed findings regarding theta’s role in retrieval processes.

We used the same decomposition approach to estimate oscillatory power during a 750 ms epoch between 800 ms and 50 ms prior to recall onset. The theta increase observed during encoding is even more pronounced at retrieval; it even shows up in traditional power decompositions like Morlet wavelets and IRASA’s mixed spectrum (see Figure 3A,B). Positive theta during retrieval has previously been reported from intracianial electrodes in the right temporal pole (Burke, Sharan, et al., 2014), but these findings were called into question by subsequent studies with far greater statistical power showing either decreases or no significant effect (Solomon, Lega, et al., 2019; Weidemann et al., 2019). We note that most prior work averaged wavelet power over the traditional theta band from either 3 or 4 Hz to 8 Hz, which blends together the positive and negative effects shown in Figure 3. Assessing a more continuous spectrum instead of averaging within bands, we recover a strong positive effect at low frequencies. So, oscillatory power obtained with IRASA for the memory retrieval contrast matches the results obtained using traditional wavelet power, though we do obtain somewhat better resolution that reveals multiple distinct components in the high theta/alpha range.

**Figure 3:**
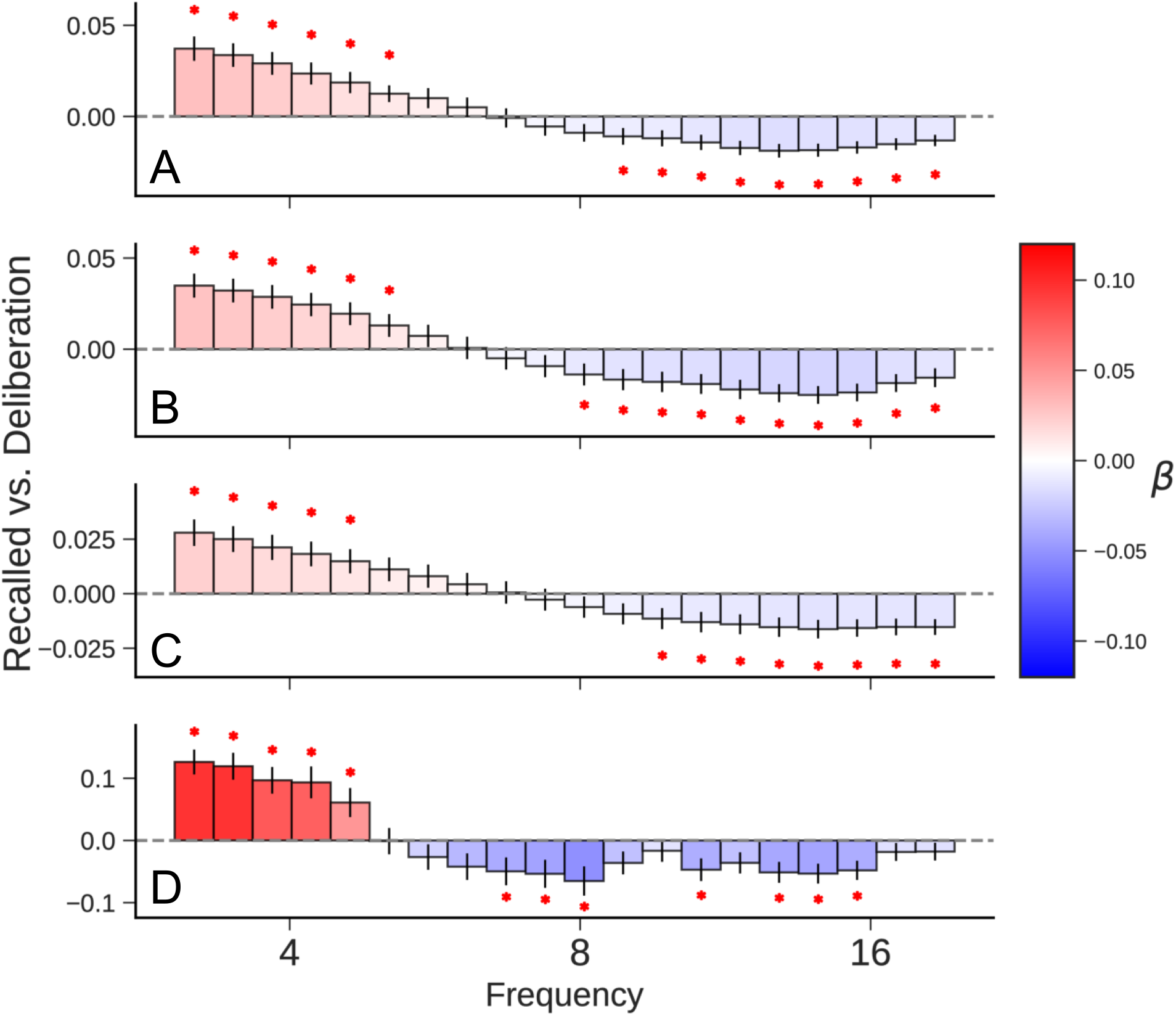
Memory Retrieval Contrast. Comparison of successful retrieval events and matched baseline deliberation events which we treat as “failed recall”. Power at logarithmically spaced frequencies were computed for the 750ms preceding recall vocalization. Values represent fixed effect coefficients for the effect of successful recall on spectral power. A red asterisk indicates that the effect size is significant after correcting for multiple comparisons (*p* < .05). Subplots show **A)** wavelet power, **B)** IRASA mixed power, **C)** IRASA fractal power, and **D)** IRASA oscillatory power.

Herweg et al (2020) proposed that choice of referencing scheme may potentially contribute to the apparent inconsistencies between non-invasive and invasive analyses of theta’s role in memory. Scalp EEG and MEG studies often report positive theta correlates of memory encoding and retrieval (Klimesch, Doppelmayr, Russegger, & Pachinger, 1996; Hanslmayr et al., 2011; Kaplan et al., 2012; Fellner, Bäauml, & Hanslmayr, 2013; Staudigl & Hanslmayr, 2013; Backus, Schoffelen, Szebényi, Hanslmayr, & Doeller, 2016) whereas many highly powered intracranial studies have failed to show these effects. The two most commonly employed - and practically distinct - methods of voltage referencing are the bipolar reference and the average reference. In a bipolar scheme, the potential difference is calculated between pairs of neighboring electrodes. This is effectively a spatial filter, as any signal shared by both electrodes will be eliminated by the differencing operation. An average reference is more sensitive to global changes in field potentials; it is calculated by averaging the potential measured at all electrodes, and subtracting that average from each one. Herweg et al (2020) observe that increases in theta power reported in scalp EEG and MEG studies with average referencing often exhibit a broad topography across the scalp, centered around frontal electrodes, and suggest that these effects might have been attenuated in intracranial studies that frequently used bipolar referencing schemes. This is because bipolar referencing acts as a spatial high-pass filter, attenuating theta effects that occur synchronously across neighboring electrodes. Comparing the memory-related power changes measured with each referencing scheme did not reveal any reliable differences (see Figure 4). An FDR-corrected paired t-test comparing bipolar to average reference (for 126 patients with monopolar recordings) did not identify significant differences between the oscillatory power estimates for the two referencing schemes at any of the frequencies of interest.

**Figure 4:**
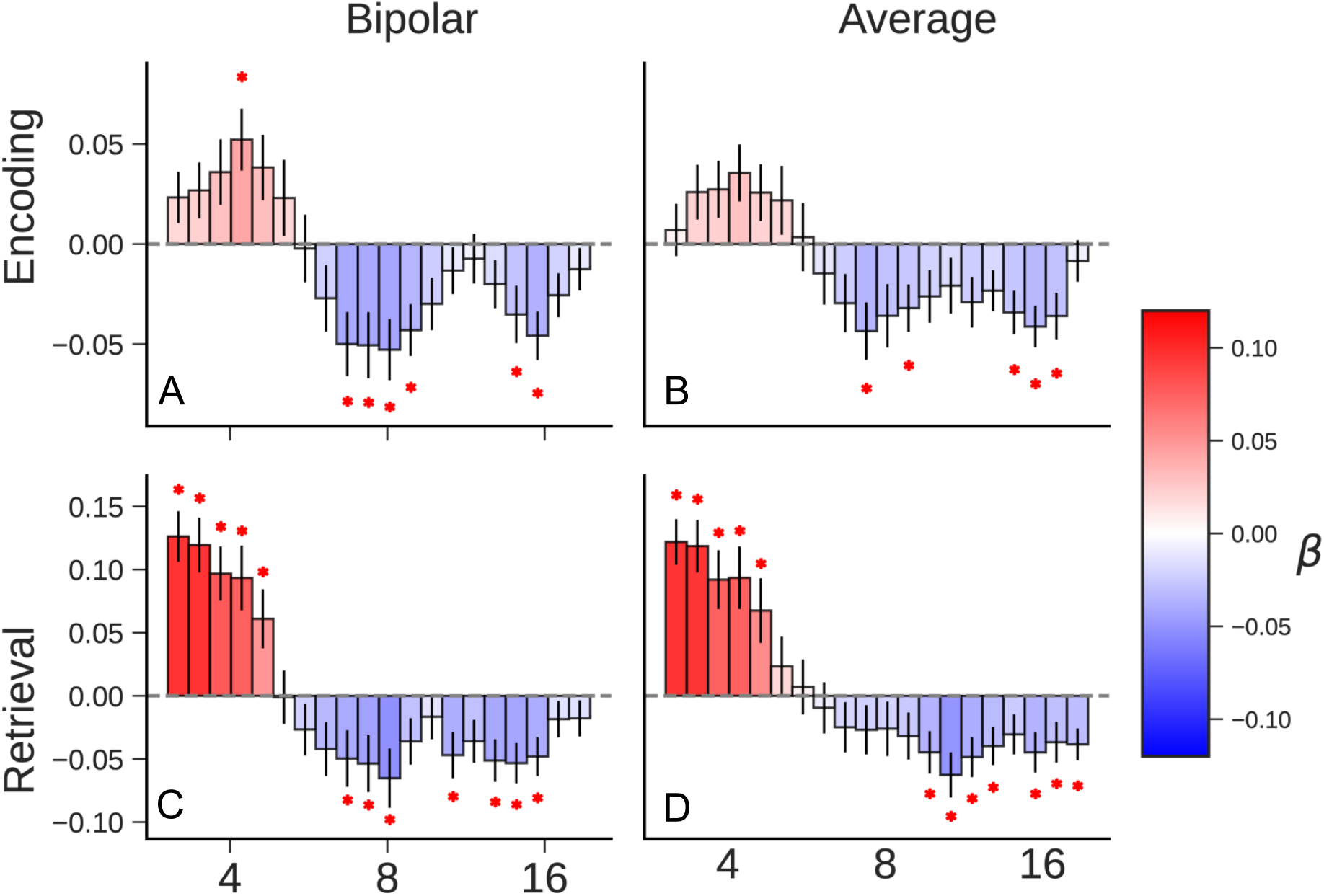
Comparison of referencing schemes. Average effect sizes for successful vs unsuccessful memory contrasts **A)** with bipolar reference at encoding, **B)** with average reference at encoding, **C)** with bipolar reference at retrieval, and **D)** with average reference at retrieval. The isolated oscillatory power spectra did not differ significantly based on the spatial filter applied to the data.

## 4 Discussion

We sought to resolve long-standing controversies regarding the role of hippocampal theta in learning and memory. To do so, we reanalyzed a large dataset of human hippocampal activity recorded as neurosurgical patients performed multiple trials of a verbal delayed free recall task. Our dataset comprised 797 hippocampal recordings across 162 patients. Whereas previous research found inconsistent theta correlates of successful encoding and recall, we find narrow-band 4-Hz oscillations to consistently increase during successful encoding (Figure 2) and preceding spontaneous free recall (as compared with matched deliberation periods, Figure 3). Further, we show that increases in theta activity appear similarly whether measured using a local spatial filter (bipolar referencing) or a more global filter (referencing to the average of all electrodes, see Figure 4).

Although many studies report theta-correlates of memory in broader memory regions, only a few studies specifically isolate hippocampal signals. Fell et al. (2011) analyzed hippocampal theta during memory encoding in a continuous recognition procedure. Analyzing ~100 hippocampal electrodes they found a significant interaction between pre- and post-stimulus presentation changes in theta power, with significant prestimulus theta increases predicting subsequent recognition. During the post-stimulus item encoding period they found a modest *decrease* in theta (*p* ~ 0.10) for subsequently recognized items. Sederberg, Schulze-Bonhage, Madsen, Bromfield, McCarthy, et al. (2007) analyzed hippocampal subsequent memory effects in a delayed free recall task. Their study, which included 186 hippocampal recordings detected reliable high-frequency increases during successful encoding, but they failed to observe reliable theta effects. In a much larger analysis of hippocampal recordings in delayed recall (401 hippocampal electrodes) Long, Burke, and Kahana (2014) observed negative theta SMEs during successful encoding, and null-effects in the theta band during successful retrieval (Burke, Long, et al., 2014). Lega, Jacobs, and Kahana (2012) examined 237 hippocampal recordings during delayed free recall (as in Sederberg et al., 2007). Recognizing the possibility that spectral measures confound broadband and narrowband (oscillatory) effects, Lega applied an oscillation detection algorithm (Caplan et al., 2003) to filter for electrodes that exhibited narrowband oscillations in each of several frequency bands. Analyzing these channels revealed both positive and negative theta effects at different electrodes. Although Lega’s study revealed striking positive theta effects at specific electrodes, it found an even larger number of hippocampal recordings that exhibited narrow-band decreases, thus offering a potential explanation for the negative and null-results described above.

Standard methods used to analyze spectral EEG power (such as wavelets, multi-tapers and windowed FFTs) mix narrow-band and broad-band signals, leaving open the possibility that a negative broadband effect can mask a positive narrowband effect, and vice-versa. When analyzed in this manner, our study replicated a number of previously published studies in showing *decreased* hippocampal theta power during the encoding of subsequently forgotten items (e.g., Burke et al., 2014; Solomon et al., 2019). By using irregular-resampling auto-spectral (IRASA), however, we revealed a positive relation between 4-Hz theta and successful memory encoding that tends to be obscured by a large negative relation between broadband power and encoding success.

Although we used IRASA to isolate narrow-band power, a number of other methods have been developed to address this problem, usually by modelling the 1/*f* background and considering deviations or residuals to be true narrowband, synchronous oscillations. The Better Oscillation Detection Method (Caplan, Madsen, Raghavachari, & Kahana, 2001) characterizes oscillations by measuring when narrowband power exceeds a power threshold above the fitted 1/*f* spectrum for a specified number of cycles at the frequency of interest; a newer method called FOOOF (Donoghue et al., 2020) identifies oscillations by assuming they are Gaussians superposed on top of a 1/*f* distribution and selecting oscillatory peaks through an iterative fitting algorithm. We expect that using these related methods would lead to similar results regarding the increase in theta with successful memory encoding.

Comparing the period immediately preceding correct recall of a studied item with matched deliberation intervals revealed that while low-frequency (4 Hz) theta increased, higher-theta band power decreased. In this case, separating narrow- and broad-band power did not prove necessary to uncover the positive correlation between theta activity and successful recall. Finally, Herweg et al hypothesized that bipolar referencing may obscure theta increases by filtering out activity correlated across multiple neighboring electrodes. Our comparison of bipolar and average references reveals clear theta increases irrespective of referencing scheme.

This study does not discuss, nor directly account for, differences in epilepsy etiologies across patients or in epileptic activity across trials. While these and other clinical factors are outside the scope of this paper, as they do not bear directly on our hypotheses, they may bear on the study of memory in patients with epilepsy more generally. (Camarillo-Rodriguez et al., 2022; Quon et al., 2021).

Hanslmayr et al. (2012) and Hanslmayr, Staresina, and Bowman (2016) propose that broadband and narrowband spectral activity have distinct and complementary roles in memory encoding: broad low-frequency desynchronization across the brain supports increased representation of information content, while narrowband theta power increases reflect the hippocampus organizing and encoding that information. Our data are consistent with this theory, as we observe simultaneous decreases in broadband, fractal power and increases in narrowband, oscillatory theta power during memory encoding. Numerous other computational models of memory, mostly informed by studies in rodents, assign a prominent role for theta in both memory formation and retrieval. Our decomposition of narrowband and broadband components of human hippocampal field potentials reveals increases in narrow-band theta during both successful encoding and retrieval, supporting the applicability of these models to human episodic memory.

## Acknowledgements

The authors would like to acknowledge our funding source for this work, NIH/NINDS Grant U01 NS1113198. Many thanks to Daniel Schonhaut, Josh Jacobs, John Sakon, David Halpern, and Noa Herz for helpful comments on previous versions of this manuscript.

## References

Avants, B. B., Epstein, C. L., Grossman, M., & Gee, J. C. (2008). Symmetric diffeomorphic image registration with cross-correlation: evaluating automated labeling of elderly and neurodegenerative brain. Medical Image Analysis, 12 (1), 26–41.

Backus, A. R., Schoffelen, J.-M., Szebényi, S., Hanslmayr, S., & Doeller, C. F. (2016, February). Hippocampal-prefrontal theta oscillations support memory integration. Current Biology, 26 (4), 450–457.

Bates, D., Kliegl, R., Vasishth, S., & Baayen, H. (2018). Parsimonious mixed models. arXiv (arXiv:1506.04967).

Bates, D., Mächler, M., Bolker, B., & Walker, S. (2015). Fitting linear mixed-effects models using lme4. Journal of Statistical Software, 67 (1), 1–48. doi: 10.18637/jss.v067.i01

Burke, J. F., Long, N. M., Zaghloul, K. A., Sharan, A. D., Sperling, M. R., & Kahana, M. J. (2014). Human intracranial high-frequency activity maps episodic memory formation in space and time. NeuroImage, 85, 834–843. doi: 10.1016/j.neuroimage.2013.06.067

Burke, J. F., Ramayya, A. G., & Kahana, M. J. (2015). Human intracranial high-frequency activity during memory processing: Neural oscillations or stochastic volatility? Current Opinion in Neurobiology, 31, 104–110. doi: https://doi.org/10.1016/j.conb.2014.09.003

Burke, J. F., Sharan, A. D., Sperling, M. R., Ramayya, A. G., Evans, J. J., Healey, M. K.,… Kahana, M. J. (2014). Theta and high–frequency activity mark spontaneous recall of episodic memories. Journal of Neuroscience, 34 (34), 11355–11365. doi: 10.1523/JNEUROSCI.2654-13.2014

Buzsaki, G., & Moser, E. (2013). Memory, navigation and theta rhythm in the hippocampal-entorhinal system. Nature Neuroscience, 16 (2), 130–138.

Camarillo-Rodriguez, L., Leenen, I., Waldman, Z., Serruya, M., Wanda, P. A., Herweg, N. A.,… Sperling, M. R. (2022). Temporal lobe interictal spikes disrupt encoding and retrieval of verbal memory: A subregion analysis. Epilepsia, 1–13. doi: https://doi.org/10.1111/epi.17334

Caplan, J. B., Madsen, J. R., Raghavachari, S., & Kahana, M. J. (2001). Distinct patterns of brain oscillations underlie two basic parameters of human maze learning. Journal of Neurophysiology, 86, 368–380.

Caplan, J. B., Madsen, J. R., Schulze-Bonhage, A., Aschenbrenner-Scheibe, R., Newman, E. L., & Kahana, M. J. (2003). Human theta oscillations related to sensorimotor integration and spatial learning. Journal of Neuroscience, 23 (11), 4726–4736.

Donoghue, T., Haller, M., Peterson, E. J., Varma, P., Sebastian, P., Gao, R.,… Voytek, B. (2020, November). Parameterizing neural power spectra into periodic and aperiodic components. Nature Neuroscience, 23, 1655–1665.

Ekstrom, A. D., Caplan, J., Ho, E., Shattuck, K., Fried, I., & Kahana, M. (2005). Human hippocampal theta activity during virtual navigation. Hippocampus, 15, 881–889.

Ezzyat, Y., Kragel, J. E., Burke, J. F., Levy, D. F., Lyalenko, A., Wanda, P. A.,… Kahana, M. J. (2017). Direct brain stimulation modulates encoding states and memory performance in humans. Current Biology, 27 (9), 1251–1258. doi: 10.1016/j.cub.2017.03.028

Ezzyat, Y., Wanda, P., Levy, D. F., Kadel, A., Aka, A., Pedisich, I.,… Kahana, M. J. (2018). Closed-loop stimulation of temporal cortex rescues functional networks and improves memory. Nature Communications, 9 (1), 365. doi: 10.1038/s41467-017-02753-0

Fell, J., Ludowig, E., Staresina, B., Wagner, T., Kranz, T., Elger, C. E., & Axmacher, N. (2011). Medial temporal theta/alpha power enhancement precedes successful memory encoding: evidence based on intracranial eeg. Journal of Neuroscience, 31 (14), 5392–5397.

Fellner, M.-C., Bäuml, K.-H. T., & Hanslmayr, S. (2013). Brain oscillatory subsequent memory effects differ in power and long-range synchronization between semantic and survival processing. NeuroImage, 79, 361–370.

Fellner, M.-C., Gollwitzer, S., Rampp, S., Kreiselmeyr, G., Bush, D., Diehl, B.,… Hanslmayr, S. (2019). Spectral fingerprints or spectral tilt? Evidence for distinct oscillatory signatures of memory formation. PLoS Biology, 17 (7), e3000403. doi: 10.1371/journal.pbio.3000403

Hanslmayr, S., Staresina, B. P., & Bowman, H. (2016, January). Oscillations and episodic memory: Addressing the synchronization/desynchronization conundrum. Trends In Neurosciences, 39 (1), 16–25.

Hanslmayr, S., Staudigl, T., & Fellner, M. (2012). Oscillatory power decreases and long-term memory: the information via desynchronization hypothesis. Frontiers in Human Neuroscience, 6 (74).

Hanslmayr, S., Volberg, G., Wimber, M., Raabe, M., Greenlee, M. W., & Bäumel, K. H. T. (2011). The relationship between brain oscillations and bold signal during memory formation: A combined eeg-fmri study. Journal of Neuroscience, 31 (44), 15674–15680.

Herweg, N. A., Solomon, E. A., & Kahana, M. J. (2020). Theta oscillations in human memory. Trends in Cognitive Science, 24 (3), 208–227. doi: https://doi.org/10.1016/j.tics.2019.12.006

Johnson, E. L., & Knight, R. T. (2015). Intracranial recordings and human memory. Current opinion in Neurobiology, 31, 18–25.

Kahana, M. J. (2020). Computational models of memory search. Annual Review of Psychology, 71 (1), 107–138. doi: 10.1146/annurev-psych-010418-103358

Kahana, M. J., Sekuler, R., Caplan, J. B., Kirschen, M., & Madsen, J. R. (1999). Human theta oscillations exhibit task dependence during virtual maze navigation. Nature, 399, 781–784.

Kaplan, R., Doeller, C. F., Barnes, G. R., Litvak, V., Düzel, E., Bandettini, P. A., & Burgess, N. (2012). Movement-related theta rhythm in humans: Coordinating self-directed hippocampal learning. PLoS biology, 10 (2), e1001267.

Klimesch, W., Doppelmayr, M., Russegger, H., & Pachinger, T. (1996). Theta band power in the human scalp EEG and the encoding of new information. NeuroReport, 7, 1235–1240.

Knierim, J., Kudrimoti, H., & McNaughton, B. (1995). Place cells, head direction cells, and the learning of landmark stability. Journal of Neuroscience, 15 (3), 1648–1659.

Lega, B., Jacobs, J., & Kahana, M. (2012). Human hippocampal theta oscillations and the formation of episodic memories. Hippocampus, 22 (4), 748–761. doi: https://doi.org/10.1002/hipo.20937

Lisman, J., Jensen, O., & Kahana, M. J. (2001). Toward a physiologic explanation of behavioral data on human memory. In C. Hölscher (Ed.), Neuronal mechanisms of memory formation (pp. 195–223). Cambridge: Cambridge University Press.

Lisman, J. E., & Jensen, O. (2013). The theta-gamma neural code. Neuron, 77 (6), 1002–1016.

Long, N. M., Burke, J. F., & Kahana, M. J. (2014). Subsequent memory effect in intracranial and scalp EEG. NeuroImage, 84, 488–494. doi: 10.1016/j.neuroimage.2013.08.052

Long, N. M., Sperling, M. R., Worrell, G. A., Davis, K. A., Gross, R. E., Lega, B. C.,… Kahana, M. J. (2017). Contextually mediated spontaneous retrieval is specific to the hippocampus. Current Biology, 27 (7), 1074–1079. doi: 10.1016/j.cub.2017.02.054

Matuschek, H., Kliegl, R., Vasishth, S., Baayen, H., & Bates, D. (2017, June). Balancing type i error and power in linear mixed models. Journal of Memory and Language, 94, 305–315.

McNaughton, B. L., Barnes, C. A., & O’Keefe, J. (1983). The contributions of position, direction, and velocity to single unit activity in the hippocampus of freely-moving rats. Experimental Brain Research, 52 (1), 41–49.

Miller, K. J., Honey, C. J., Hermes, D., Rao, R. P., den Nijs, M., & Ojemann, J. G. (2014, January). Broadband changes in the cortical surface potential track activation of functionally diverse neuronal populations. NeuroImage, 85, 711–720.

O’Keefe, J., & Dostrovsky, J. (1971). The hippocampus as a spatial map: Preliminary evidence from unit activity in the freely-moving rat. Brain Research, 34, 171–175.

Phan, T. D., Wachter, J. A., Solomon, E., & Kahana, M. J. (2019). Multivariate stochastic volatility modeling of neural data. eLife, 8, e42950.

Quon, R. J., Camp, E. J., Meisenhelter, S., Song, Y., Steimel, S. A., Testorf, M. E.,… Jobst, B. C. (2021). Features of intracranial interictal epileptiform discharges associated with memory encoding. Epilepsia.

Schacter, D. L. (1977). Eeg theta waves and psychological phenomena: A review and analysis. Biological Psychology, 5 (1), 47–82.

Scoville, W. B., & Milner, B. (1957). Loss of recent memory after bilateral hippocampal lesions. Journal of Neurology, Neurosurgreery, and Psychiatry, 20, 11–21.

Sederberg, P. B., Kahana, M. J., Howard, M. W., Donner, E. J., & Madsen, J. R. (2003). Theta and gamma oscillations during encoding predict subsequent recall. Journal of Neuroscience, 23 (34), 10809–10814. doi: 10.1523/JNEUROSCI.23-34-10809.2003

Sederberg, P. B., Schulze-Bonhage, A., Madsen, J. R., Bromfield, E. B., Litt, B., Brandt, A., & Kahana, M. J. (2007). Gamma oscillations distinguish true from false memories. Psychological Science, 18 (11), 927–932. doi: 10.1111/j.1467-9280.2007.02003.x

Sederberg, P. B., Schulze-Bonhage, A., Madsen, J. R., Bromfield, E. B., McCarthy, D. C., Brandt, A.,… Kahana, M. J. (2007). Hippocampal and neocortical gamma oscillations predict memory formation in humans. Cerebral Cortex, 17 (5), 1190–1196. doi: 10.1093/cercor/bhl030

Solomon, E. A., Lega, B. C., Sperling, M. R., & Kahana, M. J. (2019). Hippocampal theta codes for distances in semantic and temporal spaces. Proceedings of the National Academy of Sciences, 116 (48), 24343–24352. doi: https://doi.org/10.1073/pnas.1906729116

Solomon, E. A., Stein, J. M., Das, S., Gorniak, R., Sperling, M. R., Worrell, G.,… Kahana, M. J. (2019). Dynamic theta networks within the human medial temporal lobe support episodic encoding and retrieval. Current Biology, 29 (7), 1100–1111. doi: 10.1016/j.cub.2019.02.020

Squire, L. R., Knowlton, B., & Musen, G. (1993). The structure and organization of memory. Annual Review of Psychology, 44, 453–495.

Staudigl, T., & Hanslmayr, S. (2013). Theta oscillations at encoding mediate the context-dependent nature of human episodic memory. Current Biology, 23 (12), 1101–1106. doi: https://doi.org/10.1016/j.cub.2013.04.074

Voytek, B., & Knight, R. T. (2015, June). Dynamic network communication as a unifying neural basis for cognition, development, aging, and disease. Biological Psychiatry, 77 (12), 1089–1097.

Weidemann, C. T., Kragel, J. E., Lega, B. C., Worrell, G. A., Sperling, M. R., Sharan, A. D.,… Kahana, M. J. (2019). Neural activity reveals interactions between episodic and semantic memory systems during retrieval. Journal of Experimental Psychology: General, 148 (1), 1–12. doi: 10.1037/xge0000480

Wen, H., & Liu, Z. (2016, Jan). Separating fractal and oscillatory components in the power spectrum of neurophysiological signal. Brain Topography, 29 (1), 29(1):13–26.

Yushkevich, P. A., Pluta, J. B., Wang, H., Xie, L., Ding, S.-L., Gertje, E. C.,… Wolk, D. A. (2015). Automated volumetry and regional thickness analysis of hippocampal subfields and medial temporal cortical structures in mild cognitive impairment. Human Brain Mapping, 36 (1), 258–287.

